# Divergent roles of IREG/Ferroportin transporters from the nickel hyperaccumulator *Leucocroton havanensis*

**DOI:** 10.1101/2023.08.11.552958

**Authors:** Dubiel Alfonso Gonzalez, Vanesa Sanchez Garcia de la Torre, Rolando Reyes Fernandez, Louise Barreau, Sylvain Merlot

## Abstract

In response to our ever-increasing demand for metals, phytotechnologies are being developed to limit the environmental impact of conventional metal mining. However, the development of these technologies, which rely on plant species able to tolerate and accumulate metals, is partly limited by our lack of knowledge of the underlying molecular mechanisms.

In this work, we aimed to identify genes involved in nickel hyperaccumulation in the Euphorbiaceae species *Leucocroton havanensis*. Using transcriptomic data, we identified two homologous genes, *LhavIREG1* and *LhavIREG2*, encoding divalent metal transporters of the IREG/ferroportin family. Both genes are expressed at similar levels in shoots, but LhavIREG1 shows higher expression in roots. Heterologous expression of these transporters in *A. thaliana* revealed that LhavIREG1 is localized to the plasma membrane, whereas LhavIREG2 is located at the vacuole. In addition, expression of each gene induced a significant increase in nickel tolerance. Taken together, our data suggest that LhavIREG2 is involved in nickel sequestration in vacuoles of leaf cells, whereas LhavIREG1 is mainly involved in nickel translocation from roots to shoots, but could also be involved in metal sequestration in cell walls. Our results suggest that paralogous IREG/ferroportin transporters may play complementary roles in nickel hyperaccumulation in plants.

**Highlight:** The nickel hyperaccumulator *Leucocroton havanensis* endemic to Cuba, expresses two paralogous metal transporters of the IREG/ferroportin family that play distinct but complementary roles in nickel tolerance and accumulation.

## Introduction

Since the beginning of the industrial revolution, and today accelerated by the energy transition required to reduce CO_2_ emissions, metal use intensity is increasing exponentially for strategic metals. Metal mining and refining are responsible for the dispersion of metals from primary sites, increasing the risk of contamination in the natural environment and agricultural soils (Dudka and Adriano, 1997; Li *et al*., 2014; Sonter *et al*., 2020; Vidal *et al*., 2022). Phytotechnologies based on plants capable of hyperaccumulating metals have been proposed as a solution to limit the environmental impact of metal mining, remediate contaminated soils and recycle valuable metals (Suman *et al*., 2018; DalCorso *et al*., 2019; Corzo Remigio *et al*., 2020). The development of these phytotechnologies is particularly relevant in Mediterranean and tropical regions where nickel mining can have a significant impact on rich biodiversity (van der Ent *et al*., 2015). As for other crops, a better understanding of the molecular mechanisms involved in metal tolerance and accumulation will support the development of sustainable phytotechnologies using hyperaccumulators.

Metal hyperaccumulators are plant species that are able to tolerate and accumulate enormous amounts of metal in their leaves (van der Ent *et al*., 2013). More than 700 hyperaccumulators have been identified, accumulating a wide variety of metals such as nickel, zinc or manganese. However, nickel hyperaccumulators represent the vast majority of these species (Reeves *et al*., 2018). At the molecular level, metal hyperaccumulation requires high activity of genes involved in metal chelation and transport, from metal uptake by roots to metal storage in leaves (Manara *et al*., 2020). Accordingly, molecular studies, mostly performed in hyperaccumulators from the Brassicaceae family, suggest that metal hyperaccumulation has evolved from the high and constitutive expression of genes involved in the regulation of metal homeostasis in plants (Becher *et al*., 2004; Weber *et al*., 2004; Hammond *et al*., 2006; Halimaa *et al*., 2014). For example, zinc hyperaccumulation in *Arabidopsis halleri* has evolved from the triplication of the *heavy metal Atpase 4* (*AhHMA4*) gene and mutations in cis-elements leading to high expression of this metal pump. The high activity of *AhHMA4* promotes zinc loading in the xylem and thus efficient translocation of zinc to the shoots (Hanikenne *et al*., 2008). Interestingly, similar genetic mechanisms are proposed to be at the origin of zinc hyperaccumulation in *Noccaea caerulescens*, suggesting that the high expression of *HMA4* is a convergent mechanism involved in zinc hyperaccumulation in Brassicaceae (Ó Lochlainn *et al*., 2011; Craciun *et al*., 2012). Recent studies have shown that the high expression of divalent metal ion transporters of the IREG/ferroportin family (IREG/FPN) is repeatedly associated with nickel hyperaccumulation in several plant families (Merlot *et al*., 2014; Halimaa *et al*., 2014; Meier *et al*., 2018; García de la Torre *et al*., 2021).

Plant genomes contain two distinct groups of IREG/FPN transporters (Schaaf *et al*., 2006; Taniguchi *et al*., 2015; García de la Torre *et al*., 2021). The first group, represented by *Arabidopsis thaliana* AtIREG3/MAR1, encodes plastid transporters that are likely involved in the regulation of iron homeostasis in chloroplasts and mitochondria (Conte *et al*., 2009; Kim *et al*., 2021). The second group, represented by AtIREG1/FPN1 and AtIREG2/FPN2 in *A. thaliana*, encodes vacuolar and plasma membrane transporters (Morrissey *et al*., 2009). AtIREG2/FPN2 was shown to be expressed in roots in response to iron starvation and localized to the vacuolar membrane. The *A. thaliana ireg2* mutant is more sensitive to nickel, suggesting that AtIREG2 transports nickel into vacuoles to limit metal toxicity. Accordingly, plants overexpressing *AtIREG2* are more resistant to nickel and accumulate more of this metal (Schaaf *et al*., 2006). In contrast, AtIREG1/FPN1 localizes to the plasma membrane and has been proposed to mediate the loading of cobalt into the xylem for long-distance transport to the shoot (Morrissey *et al*., 2009). However, the *ireg1* mutation further increases the nickel sensitivity associated with *ireg2*, suggesting that AtIREG1 is also capable of transporting nickel. Analysis of orthologous transporters from *Medicago truncatula* (MtFPN2) and from *rice* (OsFPN1) revealed that group 2 IREG/FPN transporters can also be found in endomembranes and Golgi respectively (Escudero *et al*., 2020; Kan *et al*., 2022).

To date, functional analyses of cellular IREG/FPN transporters associated with nickel hyperaccumulation have suggested that these transporters are involved in the storage of nickel in the vacuole of leaf cells (Merlot *et al*., 2014; García de la Torre *et al*., 2021). However, the genomes of several nickel hyperaccumulators contain more than one gene encoding for group 2 IREG/FPN transporters. Interestingly, the nickel hyperaccumulator *Leucocroton* havanensis, endemic to Cuba, expresses two genes coding for group 2 IREG/FPN transporters in leaves, whereas the related nonaccumulator species *Leucocroton havanensis* apparently expresses only one gene (Jestrow *et al*., 2012; García de la Torre *et al*., 2021). This observation raises the question of the role of homologous group 2 IREG/FPN transporters in nickel hyperaccumulation.

In this work, we have studied the function of the two group 2 IREG/FPN transporters, LhavIREG1 and LhavIREG2, identified in the nickel hyperaccumulator *Leucocroton havanensis*. Our results indicate that the two genes have distinct expression patterns in roots and shoots and localize to different cellular membranes. Heterologous expression in *A. thaliana* further suggestss that the two genes have different functions in nickel transport and may therefore play distinct but complementary roles in nickel tolerance and accumulation.

## Materials and Methods

### Leucocroton havanensis plant material

*Leucocroton havanensis* seeds were collected from a single female specimen (23°04’55.5”N 82°06’46.1”W) growing on ultramafic soil in the ecological reserve “La Coca” (Havana, Cuba). Seeds and young plantlets were grown *in vitro* on Murashige and Skoog agar medium supplemented or not with 3.2 mM NiSO_4_ for 6 weeks after germination as previously described (González and Matrella, 2013). Root and shoot samples were collected separately and washed with distilled water. After rapidly removing water with a paper towel, samples were cut into 3 mm pieces and immediately placed in RNAlater (Sigma-Aldrich). Samples were transported from Cuba to France and stored at −80°C until processing.

### Transcriptomic analyses

RNA from *L. havanensis* root and leaf samples treated or not with nickel were extracted using Tri reagent (Sigma-Aldrich). RNA sequencing, *de novo* transcriptome assembly, transcriptome annotation, read mapping and differential gene expression analysis were performed essentially as described previously (García de la Torre *et al*., 2021). The *de novo* assembly of the transcriptome (Lhav_v2), was performed using RNA-seq reads from root and shoot samples of nickel-treated plantlets. The quality of the assembly was analyzed by the TransRate v1.0.3 package using the trimmed read sequences used to assemble Lhav_v2 (Smith-Unna *et al*., 2016). The completeness of the assembled transcriptome was estimated using BUSCO (v 4.0.6) and viridiplantae_odb10 lineage dataset (Seppey *et al*., 2019). Differential gene expression analysis was performed using the edgeR bioconductor package with TMM normalization and Exact test for statistical analysis, with one repetition per sample. Genes with absolute fold change |FC| ≥ 10 and adjusted p-value ≤ 0.05 were considered as differentially expressed (DE) genes.

### Reconstruction and cloning of LhavIREG coding sequences

The coding sequence of *LhavIREG1* was predicted from transcriptome Lhav_v2 contigs #981 and #4486, and from transcriptome Lhav_v1 contigs #820 and #4646. The coding sequence of *LhavIREG2* was predicted from transcriptome Lhav_v2 contig #101 and from transcriptome Lhav_v1 contigs #170 and #14018. The coding sequence of *LhavIREG3* was predicted from transcriptome Lhav_v2 contigs #6824 and #12273, and from transcriptome Lhav_v1 contigs #8277, #9347 and #4646.

The coding sequences of *LhavIREG1* and *LhavIREG2* were amplified from leaf cDNA using high-fidelity Phusion polymerase (Thermo Scientific) with gene-specific primers containing the attB recombination sequences attB1_LhavIREG1_For*/*attB2_LhavIREG1_Rev-(no)stop and attB1_LhavIREG2_For*/*attB2_LhavIREG2_Rev-(no)stop respectively (Table S1). The PCR products were recombined into pDONOR207 using GATEWAY technology (Invitrogen) to generate pDON207-*LhavIREG1(no)stop* and pDON207-*LhavIREG2(no)stop*. We confirmed the predicted sequences of the *LhavIREG1* and *LhavIREG2* coding regions by double-strand Sanger sequencing (GATC-Eurofins).

### Phylogenetic analysis

Sequences for IREG/FPN proteins were obtained from Dicots Plaza 5.0 (Van Bel *et al*., 2022), for *Arabidopsis thaliana* AthaIREG1 (AT5G03570), AthaIREG2 (AT2G38460), AthaIREG3 (AT5G26820); *Arabidopsis lyrata* AlyrIREG1 (AL4G35730), AlyrIREG2 (AL6G13070), AlyrIREG3 (AL6G38700); *Eutrema salsugineum* EsalIREG1 (Thhalv10013299m.g), EsalIREG2 (Thhalv10016525m.g), EsalIREG3 (Thhalv10003867m.g); *Teobroma cacao* TcacIREG1 (Thecc.05G005200), TcacIREG2 (Thecc.05G005300), TcacIREG3 (Thecc.05G323900); *Cucumis melo* CmelIREG1 (MELO3C026034.2), CmelIREG3 (MELO3C004177.2); *Phaseolus vulgaris* PvulIREG1 (Phvul.010G062200), PvulIREG2 (Phvul.010G062300), PvulIREG3 (Phvul.007G201800); *Nicotiana tabacum* NtabIREG1 (Nitab4.5_0000355g0170), NtabIREG2 (Nitab4.5_0012987g0010), NtabIREG3_1 (Nitab4.5_0001338g0110), NtabIREG3_2 (Nitab4.5_0004525g0030); *Chenopodium quinoa* CquiIREG1 (AUR62006302), CquiIREG2 (AUR62026347), CquiIREG3_1 (AUR62000385), CquiIREG3_2 (AUR62006741) and *Erigeron canadensis* EcanIREG1 (ECA247G0592). The sequences of *Ricinus communis* RcomIREG1 (XP_048225718.1), RcomIREG2 (XP_048235846.1), RcomIREG3 (XP_002512519.1) and *Homo sapiens* HsapFPN (XP_047300022.1) were obtained from NCBI. The sequences of *Leucocroton havanensis* LhavIREG1, LhavIREG2 proteins were translated from the corresponding cDNA sequences (OR234317, OR234318). The sequence of LhavIREG3 was deduced from the corresponding contigs. Protein alignment was performed with CLC Genomics Workbench 22.0.1 (Qiagen) using MUSCLE v3.8.425 (Edgar, 2004). The tree was constructed using maximum likelihood phylogeny (neighbor-joining, 100 bootstrap replicates) and displayed using iTol v6 (Letunic and Bork, 2021).

### Quantification of gene expression by RT-qPCR

Total RNA was extracted with TRI reagent according to the manufacturer’s instructions (Sigma-Aldrich) and purified with RNeasy Plant Mini Kit on-column DNase I treatment (Qiagen). cDNA synthesis, quantitative PCR amplification and analysis were performed as previously described (García de la Torre *et al*., 2021). The expression of *LhavIREG1* and *LhavIREG2* was normalized to the expression of the *histidine kinase 3* gene (*LhavH3K*; Lhav_v2 contig #1232), which was chosen as a reference because of its stable expression in our transcriptomic data. Primers used for RT-qPCR experiments are given in Table S1. The relative expression of *LhavIREG1* and *LhavIREG2* was quantified according to Pfaffl (2001).

### Analysis of LhavIREG1 and LhavIREG2 activity in yeast

pDON207-*LhavIREG1stop* and pDON207-*LhavIREG2stop* were recombined with pDR195- GTW (Oomen *et al*., 2009) to generate pDR195-*LhavIREG1* and pDR195-*LhIREG2*. These constructs were transformed together with pDR195-GTW and pDR195-*AtIREG2* (Schaaf *et al*., 2006) into *Saccharomyces cerevisiae* strain Y00000H strain (BY4741; *MAT*a; *leu*2Δ; *met*15Δ; *ura3*Δ). Nickel accumulation in yeast was essentially determined as previously described (Merlot *et al*., 2014; García de la Torre *et al*., 2021). Yeast cells were grown in 50 ml of liquid SD-Ura medium supplemented with 400 µM NiCl_2_ for 30 h at 28 °C with vigorous shaking. Cells were harvested by centrifugation at 4 °C and pellets were washed twice with ice-cold [10 mM EDTA, 20 mM MES pH 5.5] and once with ice-cold ultrapure water. Yeast pellets were dried at 65°C prior to elemental analysis.

### Expression of LhavIREG1 *and* LhavIREG2 *in* Arabidopsis thaliana

pDON207-*LhavIREG1nostop* and pDON207-*LhavIREG2nostop* were recombined with the plant expression vector pMUBI83 (Merlot *et al*., 2014) to express C-terminal protein fusions with the green fluorescent protein (GFP). Both constructs pMUBI83-*LhavIREG1* and pMUBI83-*LhIREG2* were transformed into the *A. thaliana ireg2-1* mutant (Schaaf *et al*., 2006) as described previously (Merlot *et al*., 2014). For each construct, three homozygous, single locus insertion, T3 lines, expressing the fluorescent protein were selected. T3 lines together with *ireg2-1* and WT (Col) were grown *in vitro* for 10 days on Hoagland agar medium containing 10 µM Fe-HBED and supplemented with 30 or 50 µM NiCl_2_. Primary root length was measured as described previously (Merlot *et al*., 2014). To measure metal accumulation, plants were grown on Hoagland agar medium containing 10 µM Fe-HBED for 7 days and then transferred to the same medium supplemented with 50 µM NiCl_2_ for 5 days. Roots and shoots samples were prepared for elemental analyses as previously described (Merlot *et al*., 2014).

### Quantitative elemental analyses

The dry weight of yeast and plant samples was measured before mineralization in 2 ml of 70% HNO_3_ for 4 h at 120 °C. Samples were diluted with ultrapure water to a total volume of 12 ml, and metal content was measured using a 4200 MP-AES spectrophotometer (Agilent technologies) as previously described (García de la Torre *et al*., 2021).

### Confocal imaging

Roots of *ireg2-1* transgenic T_2_ lines transformed with pMUBI83-*LhavIREG1* and pMUBI83-*LhavIREG2* were imaged on a Leica SP8X inverted confocal microscope (IMAGERIE-Gif platform www.i2bc.paris-saclay.fr/bioimaging/) as previously described (Merlot et al. 2014), with laser excitation at 490 nm and collection of emitted light at 500-550nm for GFP and 600-650 nm for propidium iodide.

### Statistics

GraphPad (v7.05) and the shiny application SuperPlotsOfData (Lord *et al*., 2020) were used for ttatistical analyses and data presentation.

## Results

### Analysis of leaf and root transcriptomes of *Leucocroton havanensis*

In a previous study, we generated the leaf transcriptome of *Leucocroton havanensis* (Lhav_v1) and compared it with the leaf transcriptome of the related nonaccumulator species *Lasiocroton microphyllus* to identify genes associated with the nickel hyperaccumulation trait (García de la Torre *et al*., 2021). In this study, we wanted to deepen our knowledge of the root and shoot mechanisms involved in nickel hyperaccumulation in *L. havanensis*. We sequenced RNA from roots and shoots to generate a new transcriptome assembly for this nickel hyperaccumulator species (Lhav_v2). This assembly consists of 65936 contigs with 14320 contigs larger than 1kb (Table 1). 31936 contigs (48%) were annotated with Blast2GO and 12771 contigs (19%) with Mercator 4. We then used this transcriptome as a reference to analyze the effect of nickel on gene expression in *L. havanensis* (Figure 1, Table S2). We identified 27 contigs differentially expressed (DE) in response to nickel in roots (Figure 1B), indicating a limited effect of nickel hyperaccumulation on gene expression in this tissue. Among the 19 contigs up-regulated in response to nickel in roots, contigs encoding for seed storage proteins (MapMan4 category 19.5.1) are overrepresented (4 contigs, FDR adj. p-value = 3.1e-07). In contrast, we identified 442 DE contigs in shoots, with 170 contigs up- and 272 contigs down-regulated in response to nickel (Figure 1A). This result suggests that nickel accumulation has a more pronounced effect on the shoot transcriptome. However, we could not identify enriched functional categories among DE contigs in shoots. In particular, we did not identify anyDE contigs encoding proteins involved in metal homeostasis.

**Fig. 1.**
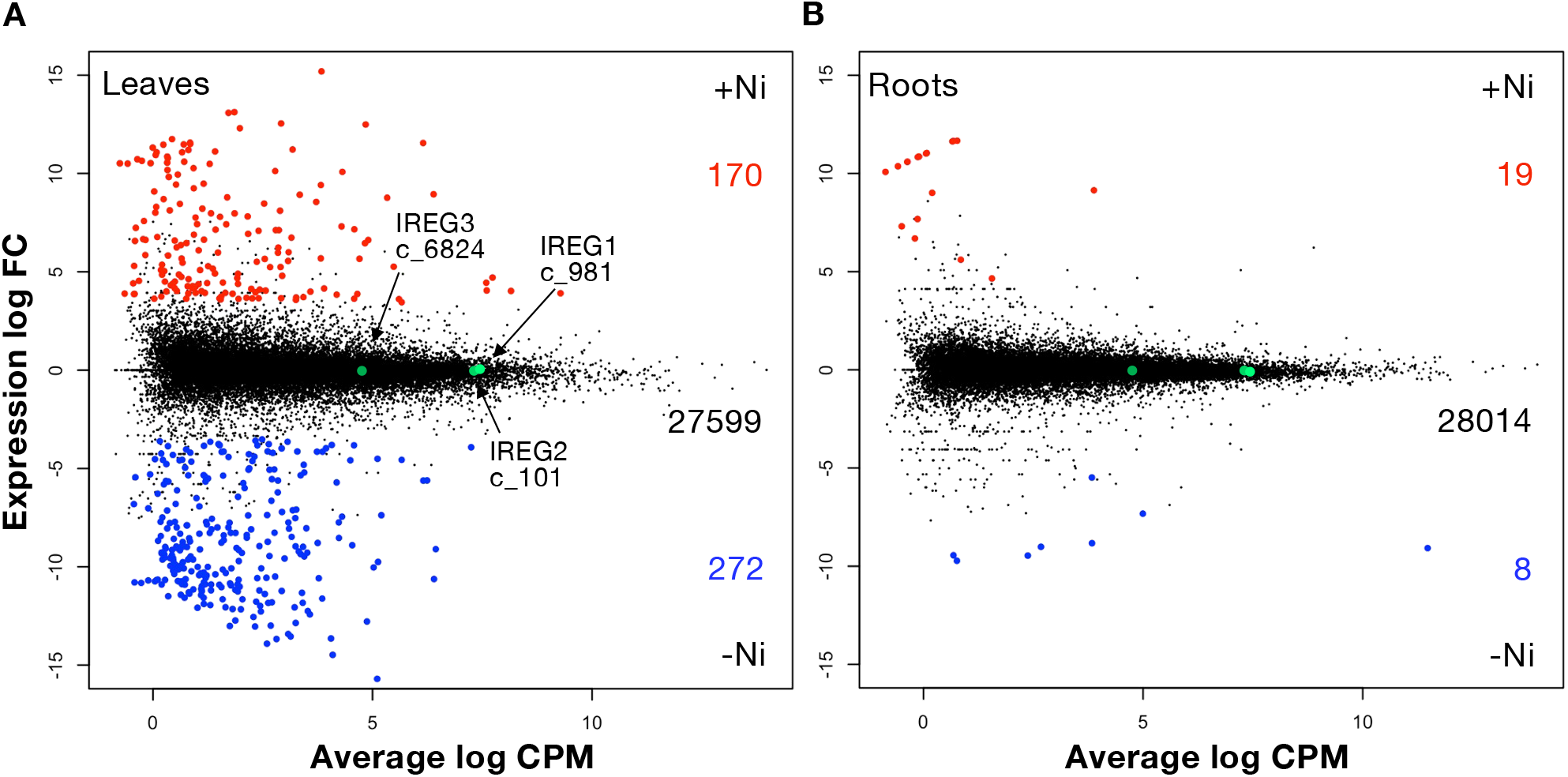
Differential gene expression analysis in *Leucocroton havanensis* in response to nickel. MA plot showing comparative analysis in shoots (A) and roots (B) in the presence (+Ni) or absence (-Ni) of nickel using the Lhav_v2 transcriptome as reference. DE contigs up-regulated in response to nickel or the absence of nickel are colored in red and blue, respectively. Non DE contigs are colored in black. The number of contigs in each category is given on the right side of the panel. The positions of *LhavIREG1* (c_981), *LhavIREG2* (c_101) and *LhavIREG3* (c_6824) are marked with green dots. FC: fold change; CPM: counts per million.

**Table 1:**
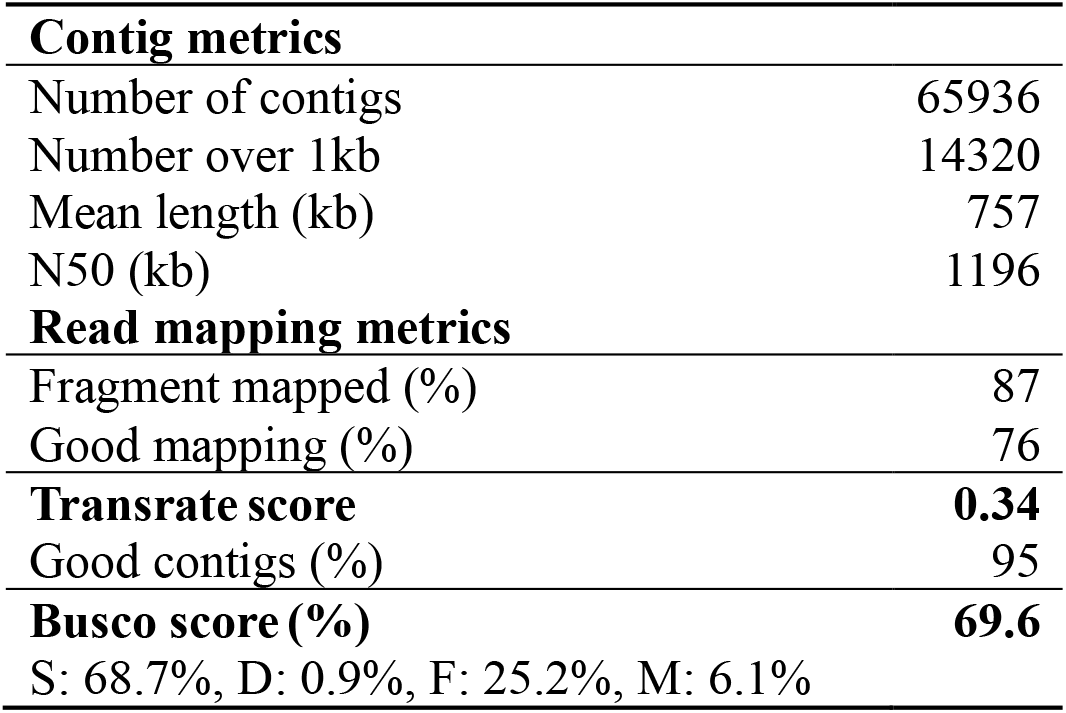
Parameters of *Leucocroton havanensis* transcriptome assembly (Lhav_v2)

### IREG/ferroportin genes in Leucocroton havanensis

IREG/FPN transporters have previously been implicated in nickel tolerance and accumulation in plants (Schaaf *et al*., 2006; Merlot *et al*., 2014; Meier *et al*., 2018; García de la Torre *et al*., 2021). Using *Leucocroton havanensis* transcriptome assemblies, we reconstructed the sequence of IREG/FPN genes expressed in this species. We identified 2 distinct transcripts coding for IREG/ferroportin transporters orthologous to *A. thaliana* AtIREG1 and AtIREG2. These transporters, subsequently named LhavIREG1 and LhavIREG2, belong to the Plaza orthologous group (OG) ORTHO05D005013, which corresponds to group 2 IREG/FPN (Figure 2; Figure S1). We identified a third transporter, LhavIREG3, orthologous to the plastidic (group 1) AtIREG3 transporter and belonging to OG ORTHO05D008050. Analysis of the phylogenetic tree obtained with plant IREG/FPN indicates that LhavIREG1 and LhavIREG2 originated from a duplication of an IREG/FPN ancestral gene in the Euphorbiaceae, independent of the duplication that occurred in the Brassicaceae and gave rise to the paralogous groups represented by AtIREG1 and AtIREG2. It is therefore not possible to infer the function of the *Leucocroton havanensis* IREG/FPN transporters from the known function of AtIREG1 and AtIREG2, which act at the plasma membrane and vacuole respectively (Schaaf *et al*., 2006; Morrissey *et al*., 2009).

**Fig. 2.**
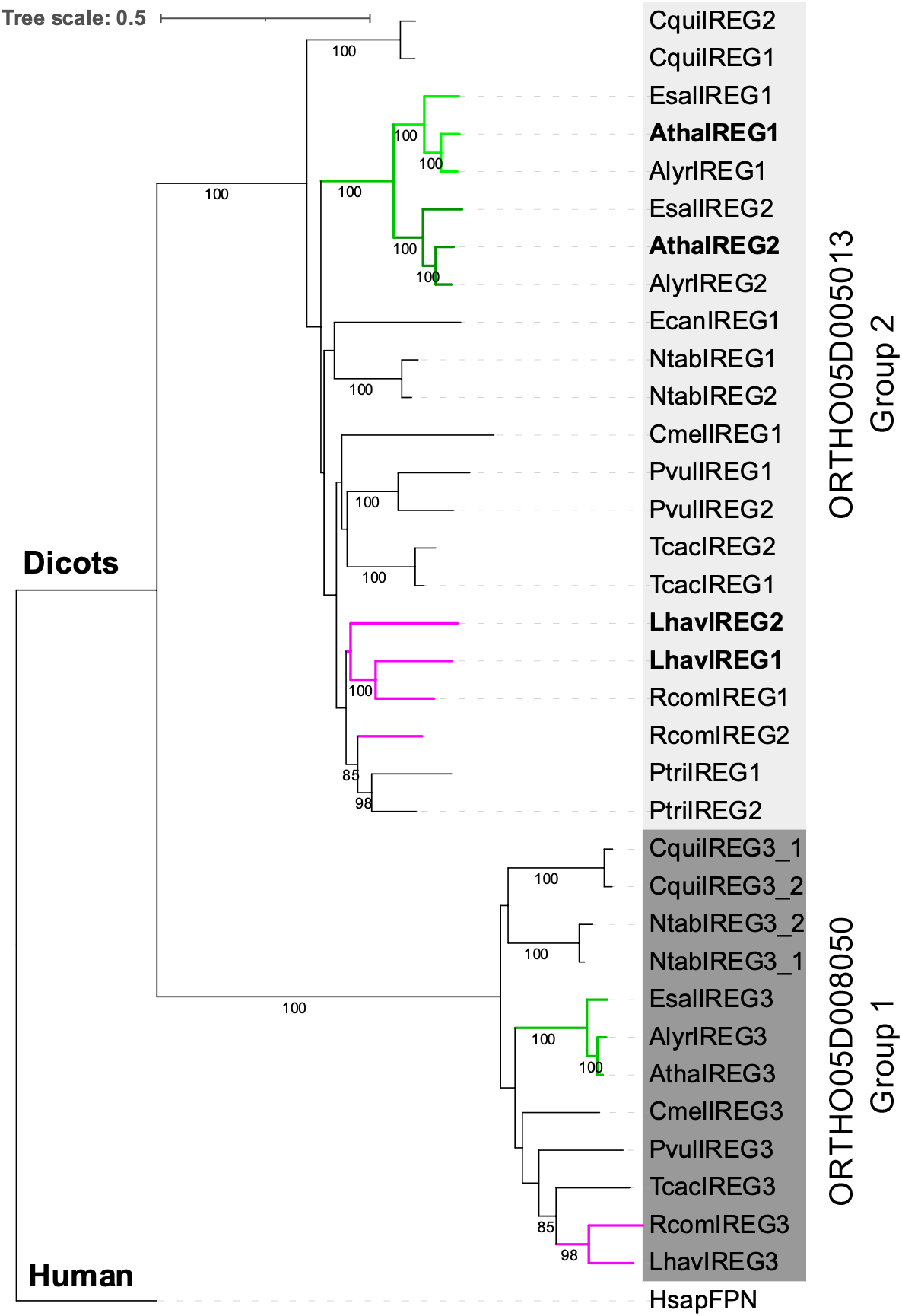
Phylogenetic tree of the IREG/FPN metal transporter family from Eudicots. This tree includes IREG/FPN proteins from *Arabidopsis thaliana* (Atha, Brassicaceae), *A. lyrata* (Alyr, Brassicaceae), *Eutrema salsugineum* (Esal, Brassicaceae), *Teobroma cacao* (Tcac, Malvaceae), *Cucumis melo* (Cmel, Cucurbitaceae), *Phaseolus vulgaris* (Pvul, Fabaceae), *Nicotiana tabacum* (Ntab, Solanaceae), *Chenopodium quinoa* (Cqui, Amaranthaceae), *Erigeron canadensis* (Ecan, Asteraceae), *Ricinus communis* (Rcom, Euphorbiaceae), *Leucocroton havanensis* (Lhav, Euphorbiaceae). *Homo sapiens* (Hsap) ferroportin was used as an outgroup. Orthogroups ORTHO05D005013 and ORTHO05D008050 from Plaza 5 are shaded in light and dark gray, respectively. Clades containing Brassicaceae (green) and Euphorbiaceae (magenta) species are highlighted.

### Analysis of LhavIREG1 and LhavIREG2 expression in Leucocroton havanensis

We analyzed the expression of *LhavIREG1 and LhavIREG2* in *L. havanensis* by RT-qPCR (Figure 3). This analysis revealed that both genes are expressed at a similar level in shoots but that *LhavIREG1* is more expressed than *LhavIREG2* in roots. These results are supported by the analysis of RNA-Seq data (Table S2). Based on RNA-Seq data, we did not observe a significative effect of nickel on the expression of IREG/FPN genes in *L. havanensis* (Figure 1). These results suggest that both genes are constitutively expressed in *L. havanensis* and that *LhavIREG1* predominantly acts in roots while both genes are likely playing a function in leaves.

**Fig. 3.**
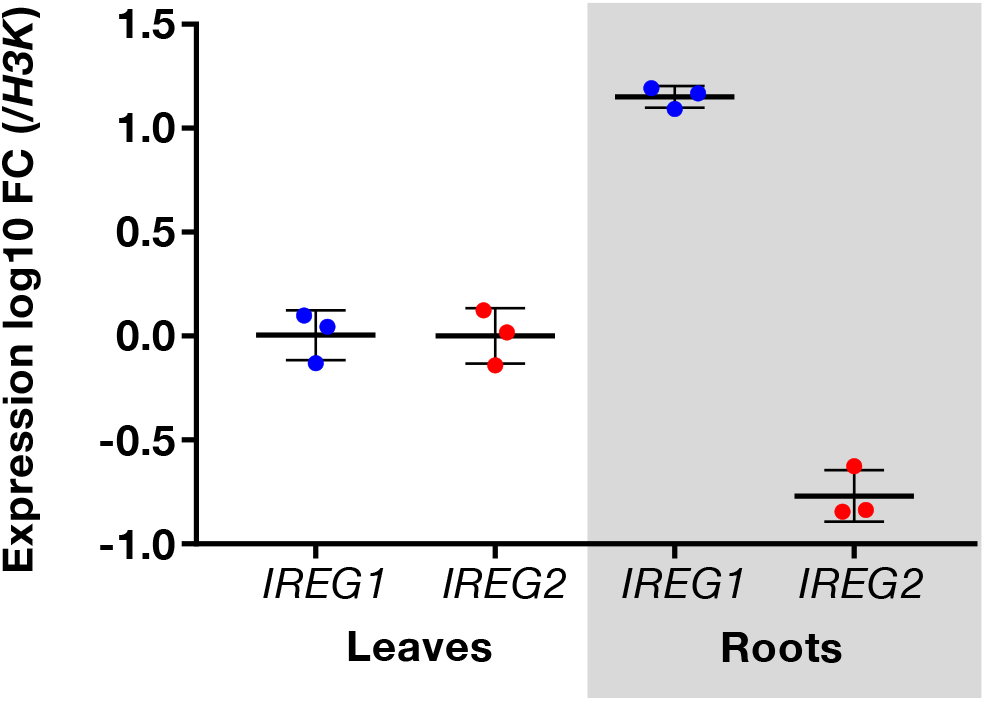
Analysis of the expression of *LhavIREG1* and *LhavIREG2* in leaves and roots. Gene expression measured by RT-qPCR is expressed as log10 fold change (FC) compared to *H3K* expression. Data are mean value ± SD (n=3 independent samples).

### LhavIREG1 and LhavIREG2 act as nickel exporters

We have previously shown that *LhavIREG2* expression in yeast increases nickel tolerance (García de la Torre *et al*., 2021). To further characterize the activity of LhavIREG2 and LhavIREG1, we measure nickel accumulation in yeast cells expressing each of these transporters (Figure 4). The expression of both LhavIREG1 and LhavIREG2 reduces nickel accumulation compared to yeast transformed with the control vector. The same result is observed with *A. thaliana* AtIREG2. These results suggest that both LhavIREG1 and LhavIREG2 transport nickel out of yeast cells, which is consistent with the conserved role of IREG/FPN as divalent metal exporters.

**Fig. 4:**
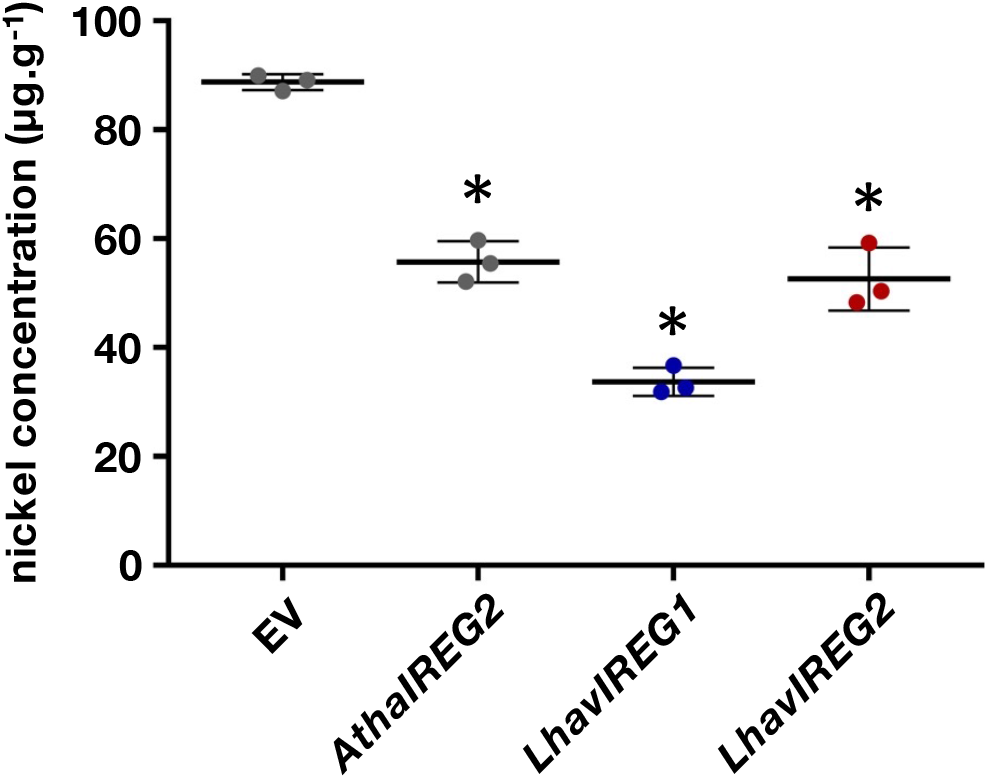
Nickel accumulation in yeast cells expressing *LhavIREG1* and *LhavIREG2.* Nickel concentration was measured by MP-AES in yeast cells expressing *LhavIREG1, LhavIREG2* and *AthaIREG2* or transformed with pDR195 (EV). For each construct, the results are mean value ± SD (n = 3; independent transformants). * *p* < 0.01 (Welch’s t-test), comparedwith EV condition

### LhavIREG1 and LhavIREG2 localize to different membranes in plant cells

To analyze the localization of LhavIREG1 and LhavIREG2 in plant cells, we fused the two transporters with the green fluorescent protein (GFP) at the C-terminal end and expressed the fusion proteins in the *A. thaliana ireg2* mutant. Root cells of the transgenic T2 lines were imaged by confocal microscopy (Figure 5). Lines expressing LhavIREG1-GFP show a thin GFP fluorescent signal delineating the periphery of the root cells (Figure 5 A, B). This signal indicates that LhavIREG1 is mainly localized to the plasma membrane. In contrast, the GFP signal associated with the expression of LhavIREG2-GFP outlines large cytoplasmic vesicles, suggesting a vacuolar localization (Figure 5 C, D). These results indicate that LhavIREG1 and LhavIREG2 localize to different membranes and therefore likely have distinct cellular functions.

**Fig. 5.**
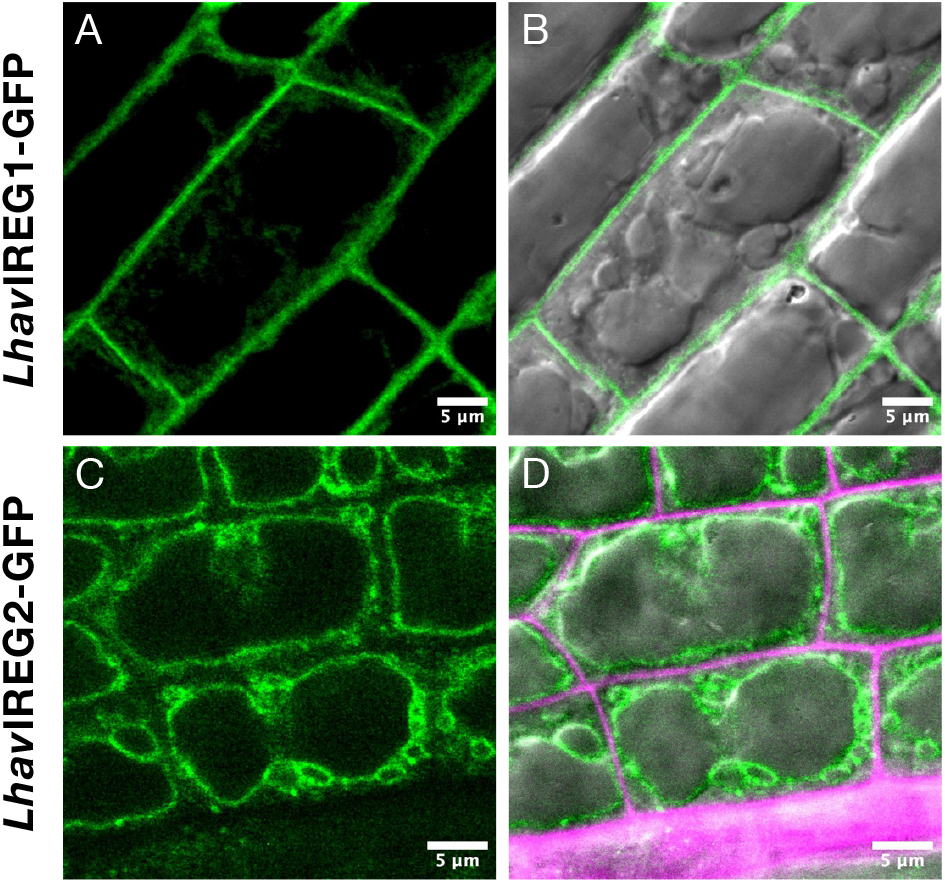
Localization of *Lhav*IREG1 and *Lhav*IREG2 in plant cells. Root cells of *A. thaliana ireg2* mutant stably expressing *Lhav*IREG1-GFP (A, B) and *Lhav*IREG2-GFP (C, D) were imaged by confocal microscopy. GFP-associated fluorescence is shown in green (A, C). Corresponding composite images with DIC images are also shown (B, D). Propidium iodine Cell walls labelled with are shown in magenta. Scale bars are 5 µm.

### Expression of LhavIREG1 and LhavIREG2 in A. thaliana differentially affects nickel sensitivity and accumulation

We used the same *ireg2* transgenic lines expressing *LhavIREG1-GFP* and *LhavIREG2-GFP* to study the effect of the ubiquitous expression of these transporters on nickel tolerance and accumulation (Figure 6). As observed in previous works (Schaaf *et al*., 2006; Merlot *et al*., 2014), the *A. thaliana ireg2* mutant is more sensitive to root growth in the presence of nickel than the wild type (Figure 6A, B, C). Expression of both *LhavIREG1-GFP* and *LhavIREG2-GFP* in *ireg2* significantly increases root growth in the presence of 30 µM nickel compared to *ireg2* or the wild-type (Figure 6A, B, C, D). In addition, transgenic lines expressing *LhavIREG1-GFP* and *LhavIREG2-GFP* do not exhibit the chlorotic phenotype observed in *ireg2* leaves in the presence of nickel. Transgenic lines expressing *LhavIREG1-GFP* show a strong tolerance to 50 µM nickel, a concentration that strongly affects the root growth of the other genotypes (Figure 6D).

**Fig. 6.**
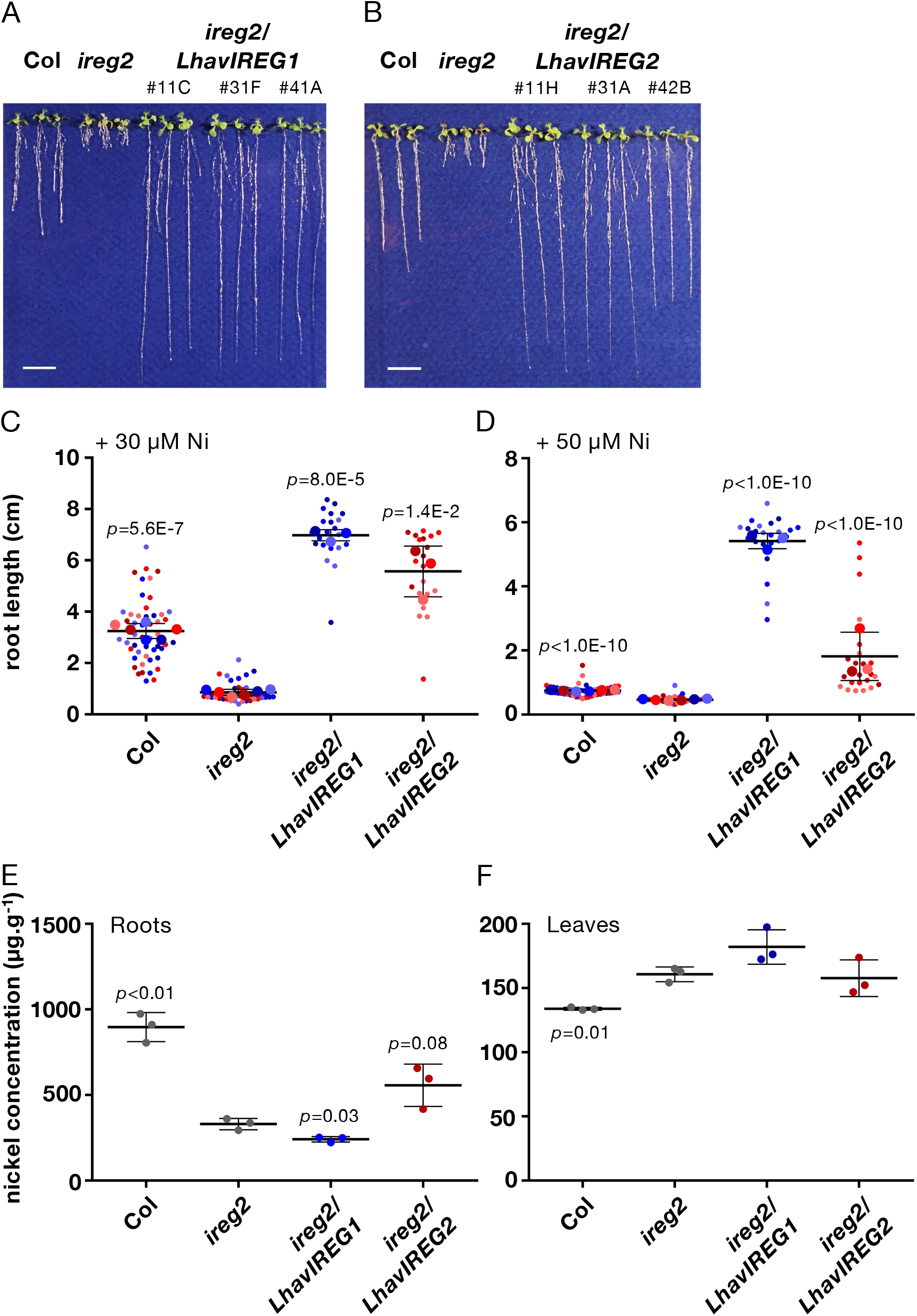
Effect of LhavIREG1 and LhavIREG2 expression on nickel tolerance and accumulation. Pictures of individual *A. thaliana ireg2* transgenic lines expressing *LhavIREG1-GFP* (A) and *LhavIREG2-GFP* (B) grown on a culture medium containing 30 µM NiCl_2_ for 10 days together with wild type (Col) and the *ireg2* mutant. Three representative plantlets were selected per line. Scale bars correspond to 1 cm. (C) Quantification of root growth of plantlets grown in the presence of 30 µM NiCl_2_ and (D) 50 µM NiCl_2_. Each small dot represents an individual measurement. Large dots represent the mean value for each individual line or experiment. Means (n=3 independent transgenic lines, n=6 for Col and *ireg2*) ± SEM are shown. *p*-values were calculated compared to *ireg2*. Nickel concentrations were measured in roots (E) and leaves (F) of the same genotypes. Data are mean value ± SD (n=3 independent transgenic lines or experiments). *p*-values were calculated relative to *ireg2*.

To further characterize these transgenic lines, we measured nickel accumulation in both roots and shoots (Figure 6E, F). The *ireg2* mutant, which is impaired in the vacuolar sequestration of nickel in roots, translocates and accumulates more nickel in shoots (Schaaf *et al*., 2006; Merlot *et al*., 2014). Expression of the vacuolar *LhavIREG2* in *ireg2* transgenic lines slightly increases nickel accumulation in roots, but does not restore nickel accumulation as in wild type (p=0.02), indicating a partial complementation of the *ireg2* phenotype. In contrast, expression of *LhavIREG1* further reduces nickel accumulation in roots compared to the *ireg2* mutant (p=0.03). In these experiments, we cannot observe a significant effect of *LhavIREG1-GFP* and *LhavIREG2-GFP* expression on nickel accumulation in leaves compared to *ireg2* (Figure 6F), but plants expressing *LhavIREG1-GFP* accumulate significantly more nickel in leaves than wild type (p=0.02). These results indicate that the ubiquitous expression of both *LhavIREG1* and *LhavIREG2* leads to a large increase in nickel tolerance, albeit through different cellular mechanisms.

## Discussion

In this work, we provided new molecular insights into the mechanisms involved in nickel hyperaccumulation in the Cuban endemic species *Leucocroton havanensis* (Jestrow *et al*., 2012; González and Matrella, 2013). We generated a novel transcriptome assembly containing genes expressed in roots and shoots in this species (Table 1). Using this assembly as a reference, we examined the expression of genes in response to nickel treatment and thus nickel hyperaccumulation (Fig. 1). Our results indicate that nickel treatment elicits a more complex response in shoots (442 DE genes) than in roots (27 DE genes). Enrichment analysis revealed that genes encoding for seed storage proteins are induced by nickel in roots. The biological significance of this response is unclear. No other functional category, including the regulation of metal homeostasis, is enriched in response to nickel. In recent years, metal transporters of the IREG/FPN family have been identified as candidate genes involved in nickel tolerance and accumulation (Schaaf *et al*., 2006; Merlot *et al*., 2014; Meier *et al*., 2018; García de la Torre *et al*., 2021). Analysis of the expression of the two genes encoding group 2 IREG/FPN transporters, *LhavIREG1* and *LhavIREG2*, indicates that both genes are significantly expressed in *L. havanensis*, but are not regulated at the transcriptional level by nickel (Fig. 1, 3). These results are in line with a model suggesting that nickel hyperaccumulation at the species level is associated with a high and constitutive expression of genes involved in metal homeostasis and transport (Verbruggen *et al*., 2009; Krämer, 2010).

We further investigated the putative function of LhavIREG1 and LhavIREG2 in nickel hyperaccumulation. Our data showed that LhavIREG1 and LhavIREG2 are both nickel exporters (Fig. 4), but localize to the plasma membrane and the vacuole, respectively (Fig. 5). The equivalent dual localization was previously observed for *Arabidopsis thaliana* AthaIREG1 and AthaIREG2 (Schaaf *et al*., 2006; Morrissey *et al*., 2009). This observation suggests that in these two distant genera, the duplication of an ancestral group 2 IREG/FPN gene resulted in two transporters that convergently evolved into a plasma membrane and a vacuolar form. The molecular determinants of the localization of the IREG/FPN transporters are not yet known. Recently, the presence of a dileucine motif [D/E]X_3-5_L[L/I] (Bonifacino and Traub, 2003; Komarova *et al*., 2012) was shown to be involved in the vacuolar localization of *A. thaliana* NRAMP3 and NRAMP4 transporters (Müdsam *et al*., 2018). At least three canonical dileucine motifs are found in the large cytoplasmic loop and C-terminal extension of LhavIREG2 but not LhavIREG1 (Fig. S1). Further studies will be required to support the role of these motifs in the vacuolar localization of LhavIREG2.

As anticipated, ectopic and constitutive expression of the vacuolar LhavIREG2 increases nickel tolerance in the *A. thaliana ireg2* mutant. Surprisingly, expression of the plasma membrane LhavIREG1 increases nickel tolerance to a greater extent (Fig. 6). Elemental analysis indicates that LhavIREG1 expression further reduces nickel accumulation in roots compared to *ireg2* and simultaneously increases accumulation in shoots. These results suggest that ectopic expression of LhavIREG1 does not drive the export of nickel from root cells, but rather promotes its translocation to the shoots. This proposed function of LhavIREG1 in metal translocation was previously proposed for *A. thaliana* IREG1 (Morrissey *et al*., 2009). *A. thaliana ireg2* plants expressing LhavIREG1 show no symptoms of nickel toxicity in leaves, suggesting nickel is accumulated in a compartment where it is less toxic. Since LhavIREG1 localizes to the plasma membrane, we propose that nickel is exported out of leaf cells and bound to the cell wall. Further experiments, including elemental imaging, would be required to support this hypothesis. Interestingly, while vacuolar sequestration of nickel is recognized as an essential function for nickel tolerance and hyperaccumulation (Schaaf *et al*., 2006; Merlot *et al*., 2014; García de la Torre *et al*., 2021), nickel is also associated with the cell wall in leaves of several hyperaccumulators (Krämer *et al*., 2000; Bidwell *et al*., 2004; van der Ent *et al*., 2019). Taken together, our data suggest that in addition to vacuolar IREG/Ferroportin transporters, plasma membrane IREG/FPN also play an important role in hyperaccumulation, mediating efficient translocation of nickel from roots to shoots and nickel export from leaf cells to limit its toxicity.

## Data and code availability

The data discussed in this publication have been deposited in NCBI’s Gene Expression Omnibus (Edgar *et al*., 2002) and are accessible through GEO Series accession number GSE237255 (https://www.ncbi.nlm.nih.gov/geo/query/acc.cgi?acc=GSE237255). Previous Leucocroton data (Lhav_v1) are available through GEO GSE116049. *LhavIREG1* and *LhavIREG2* cDNA sequences were deposited to GenBank under OR234317 and OR234318 accession numbers respectively.

## Contributions

DAG and SM: conceptualization; DAG, VSGT and SM: methodology; DAG, VSGT and SM: formal analysis; DAG, VSGT, RRF, LB and SM: investigation; DAG: resources; VSGT and SM: data curation; DAG and SM: writing - review & editing; DAG, VSGT and SM: visualization; DAG, VSGT and SM: supervision; SM: funding acquisition.

## Acknowledgement / Funding

We thank Ludivine Soubigou-Taconnat for the coordination of the *Leucocroton havanensis* RNA sequencing at the POPS platform (Orsay, France), Celeste Belloeil for help with data presentation and Sébastien Thomine for comments on the manuscript. This work was supported by the ANR funding EvoMetoNicks (ANR-13-ADAP-0004) and the MITI-CNRS Defi Enviromics GENE-4-CHEM to S.M. D.A.G and R.R.F were recipient of SCAC mobility fellowships from the French Embassy in Cuba

## Notes

### Competing Interest Statement

The authors have declared no competing interest.

